# Relative growth of invasive and indigenous tilapiine cichlid fishes in Tanzania

**DOI:** 10.1101/743740

**Authors:** SJ Bradbeer, BP Ngatunga, GF Turner, MJ Genner

## Abstract

Non-native species have been widely distributed across Africa for the enhancement of capture fisheries, but it can be unclear what benefits in terms of fisheries production the non-native species bring over native species. Here we compared the relative growth rate of sympatric populations of introduced *Oreochromis niloticus* (Nile tilapia) to indigenous *Oreochromis jipe* (Jipe tilapia) at three impoundments in Northern Tanzania. Using scale increments as a proxy for growth, we found that *O. niloticus* had an elevated growth rate relative to *O. jipe*, with the greatest *O. niloticus* growth rates being evident in the Nyumba ya Mungu reservoir. These results help explain why *O. niloticus* may be a superior competitor to native species in some circumstances. However, further introductions of this non-native species should be undertaken with caution given potential for negative ecological impacts on threatened indigenous tilapia species.

---

Invasive species are largely considered to have superior traits relative to their indigenous counterparts, enabling their establishment and success in invaded ranges. Characteristics associated with invasion success in fish include fast growth, broad environmental tolerances and high fecundity (Kolar and Lodge 2002; Moyle and Marchetti 2006). These advantageous traits have been studied alongside environmental characters of the habitat to both evaluate impacts of invasive species, as well as predict future invasions (Copp et al. 2009; Marr et al. 2017). In some circumstances, invasive species can outcompete established indigenous competitors for limited resources, such as food, breeding habitat and shelter (Bøhn et al. 2008). However, in aquatic systems while competition is often inferred based on abundance trends, or shared patterns of resource use, often there is little evidence of the relative performance of the species where they co-occur.

One indicator of fitness among sympatric species is growth. In fish, growth can be measuring using a range of methods including quantifying the deposition of calcified structures on the anterior of scale circuli (Cheung et al. 2007; Martin et al. 2012). Higher individual growth rates are considered advantageous across populations as they enable those individuals to reach reproductive age quicker, with less time spent at a more vulnerable juvenile life stage (Sutherland 1996). Furthermore, in female fish body size is directly related to egg output potential, therefore larger body sizes can enhance reproductive output (Barneche et al. 2018). Greater body size may pose an advantage for males in competition for spawning territories. Thus, comparing growth rates of species can also acts as an indication of potential relative competitiveness (Chifamba and Videler 2014).

*Oreochromis niloticus* (Nile tilapia (Linnaeus 1758)) is native to much of northern Africa, including the Nile and Niger systems (Trewavas 1983). In Tanzania, the species is naturally distributed only in the Lake Tanganyika catchment (Shechonge et al. 2019a), but due to favourable characteristics, such as potential for rapid growth and broad habitat tolerances, it has been widely distributed to non-native habitats across the country (Shechonge et al. 2019b). Such introductions have largely been deliberate, to promote capture fisheries, but it is also possible accidental escapes have occurred from aquaculture facilities in the country. Where *O. niloticus* is present in Tanzania, it typically co-occurs with indigenous tilapiine species (Bradbeer et al. 2019; Shechonge et al. 2019b). However, the fundamental ecological characteristics of populations of *O. niloticus* relative to those of native species are typically unknown, including the relative performance of fisheries-related traits such as growth rates.

Here, we report a study comparing the relative growth of introduced *O. niloticus* to indigenous *Oreochromis jipe* (Jipe tilapia (Lowe 1955)), a large bodied species endemic to the Pangani catchment that partially supports multiple artisanal fisheries in the region (Shechonge et al. 2019b). When first described, this ‘species’ was believed to represent a complex of three closely-related morphologically similar species, *O. jipe* and *O. girigan* occupying different niches within Lake Jipe and *O. pangani* within the river (Lowe 1955), but these have not been studied in depth since and have generally not been distinguished by subsequent workers and are now treated as a single species (Seegers et al. 2003; Fricke et al. 2019). We sampled fisher’s catches from three locations, namely Lake Kumba (5°1.92’ S, 38°32.88’ E), Nyumba ya Mungu reservoir (3°36.72’ S, 37°27.54’ E) and the Pangani Falls reservoir (5°20.82’ S, 38°38.7’ E) in August 2015 (Figure 1). All fishes were identified to species, individually labelled, and stored in 70% ethanol. To assess growth rates, we followed the method of Martin et al. (2012) that has been validated as a method of comparing recent growth of tilapiine cichlid fishes in experimental trials. For each specimen, three scales were removed from the area superior to the lateral line and posterior to the head. Scales were then placed onto a microscope slide, treating with glycerol and covered with a glass coverslip. Images with a superimposed scale bar were taken using a M205C microscope (Leica, Wetzlar, Germany). Image files were loaded into tpsDIG 2.2 (Rohlf 2015) and from each scale, five measurements were recorded, namely the scale total width (longest distance across the scale; Figure 2a) and four separate “increment size” measurements of the distance between the first and fifth circuli on primary radii viewed from the anterior field of the scale (Figure 2b). From these measurements we calculated a mean scale width of the individual, and the mean increment size of the individual.

**Figure 1.**
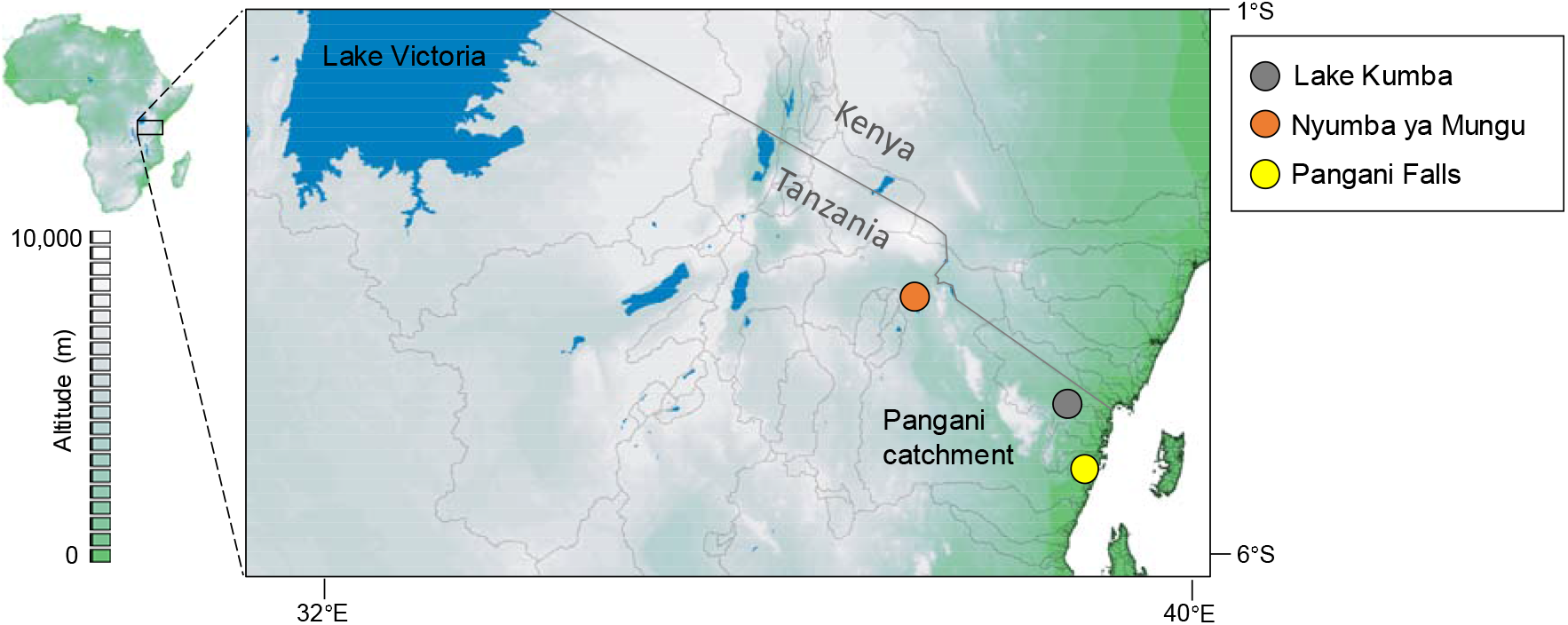
Sampling sites for sympatric Nile tilapia and Jipe tilapia in August 2015.

**Figure 2.**
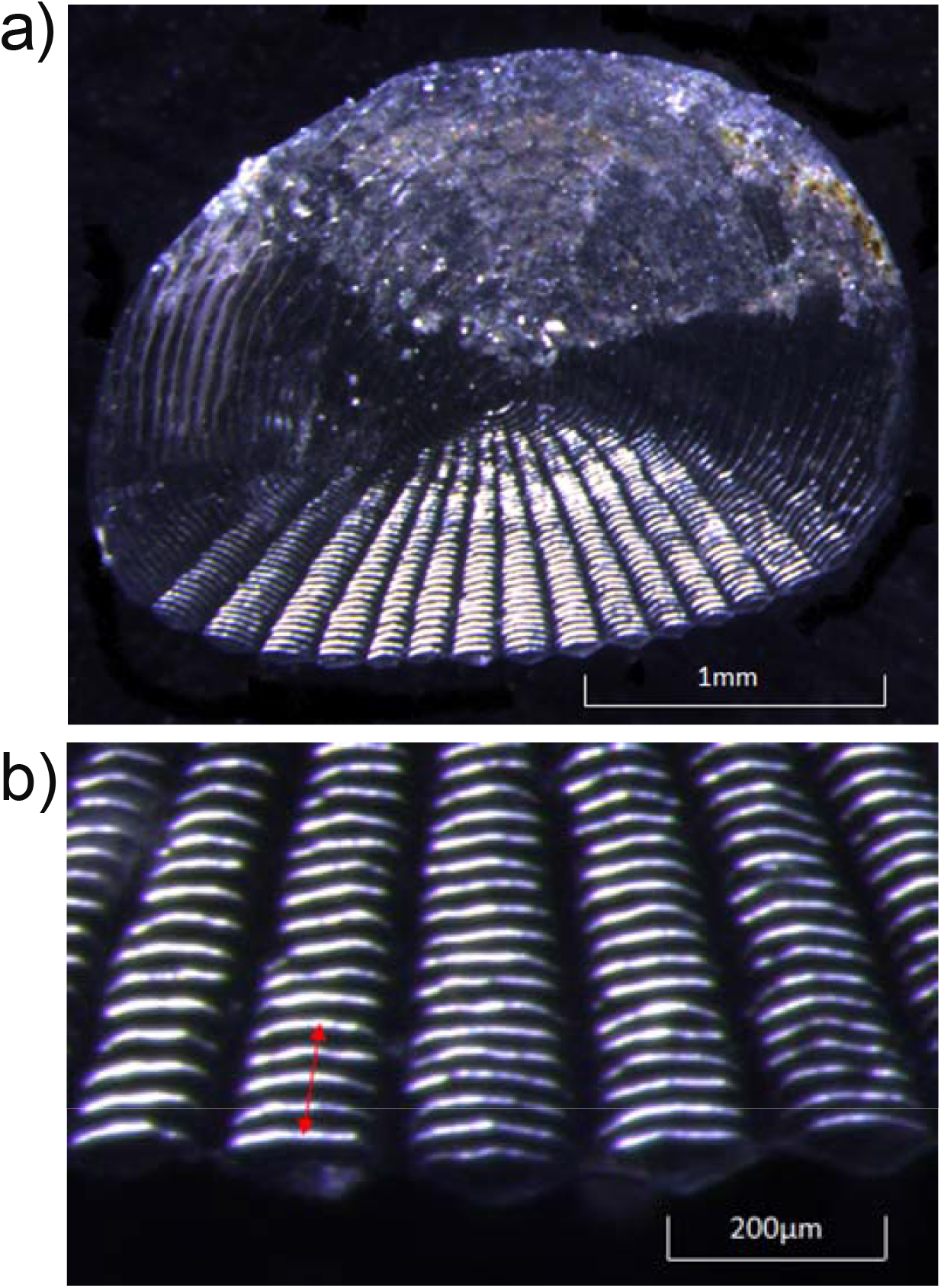
Measurements were made of a) total scale width, and b) the distance between the first and fifth circuli (indicated by arrow) on a primary radius of the same scale.

Scale total width was employed as a covariate of increment size, alongside the factors Species and Sampling Site, in an analysis of covariance in R 3.6.0 (R Core Team 2019), with size-standardised increment size (hereafter termed “relative growth”) compared using marginal means and *post-hoc* pairs tests in the emmeans package (Lenth et al. 2018). After accounting for the significant positive associations of scale total width on increment size (*F*_1,142_ = 138.53, *P* < 0.001), there were significant overall differences among sites in increment size (*F*_2,142_ = 57.55, *P* < 0.001), and *O. niloticus* had a relatively greater increment size than *O. jipe (F*_1,142_ = 30.49, *P* < 0.001). However, the extent of the difference differed significantly among locations (*F*_2,142_ = 12.72, *P* < 0.001; Figure 3). Pairwise *post-hoc* Tukey’s tests confirmed a significantly greater growth of *O. niloticus* relative to *O. jipe* at Nyumba ya Mungu (*t* = −7.303, *P* < 0.001), but no significant differences between the species at either Lake Kumba (*t* = −0.946, *P* = 0.346) or the Pangani falls (*t* = −1.427, *P* = 0.156). Pairwise *post-hoc* Tukey’s tests confirmed *O. niloticus* has a significant higher growth rate at Nyumba ya Mungu than Pangani falls (*t* = −4.710, *P* < 0.001) and Lake Kumba (*t* = −11.629, *P* < 0.001), while it also grew faster at Pangani falls than Lake Kumba (*t* = 5.625, *P* < 0.001). Pairwise *post-hoc* tests showed that *O. jipe* grew faster at Nyumba ya Mungu than Lake Kumba (*t* = −2.876, *P* = 0.013), but there were no significant differences in growth rates between the Pangani Falls population and either Nyumba ya Mungu (*t* = 0.245, *P* = 0.967) or Lake Kumba (*t* = 2.364, *P* = 0.051).

**Figure 3:**
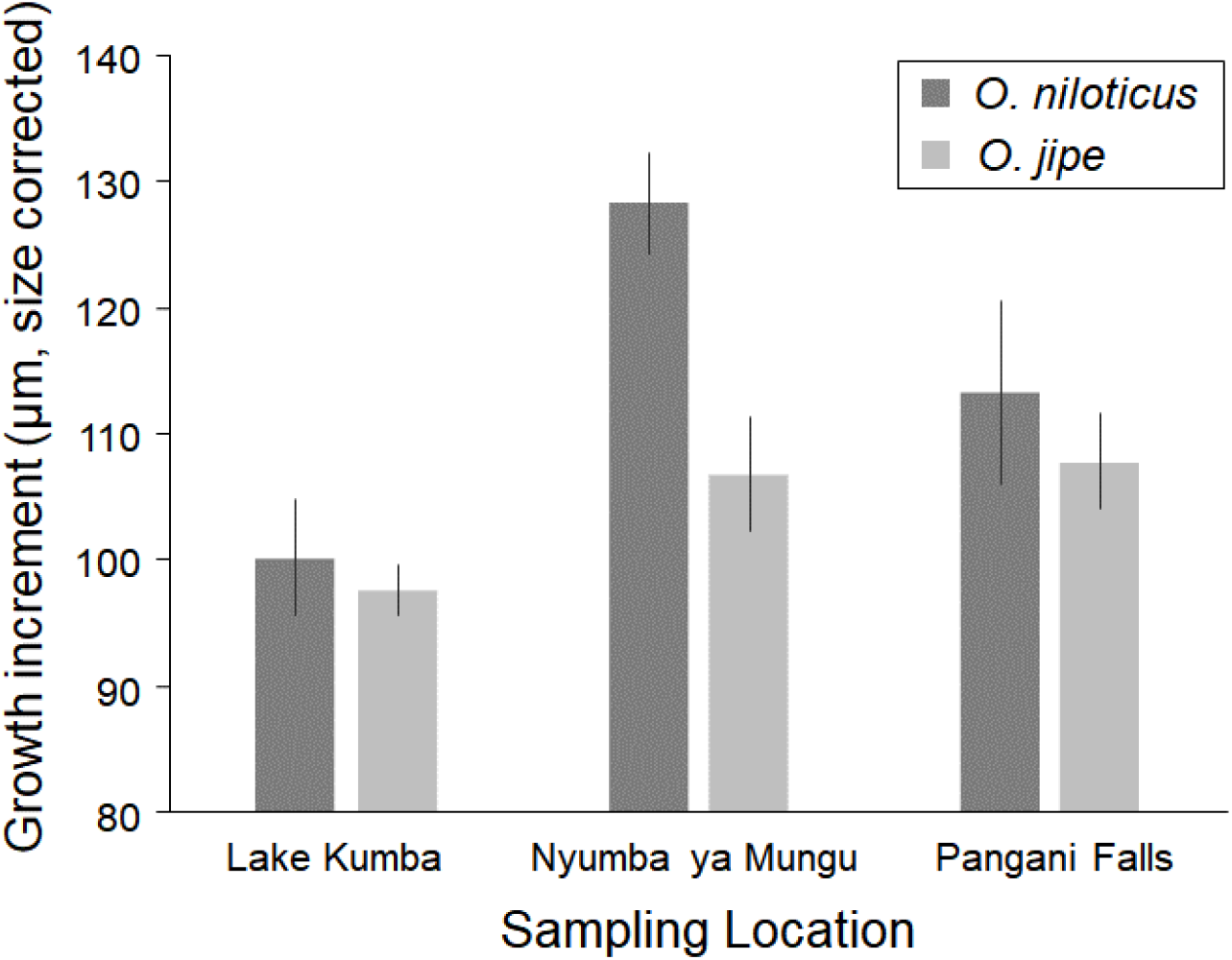
Scale growth measurements (corrected for scale width) for *Oreochromis niloticus* (dark grey) and *Oreochromis jipe* (light grey) at three sites, Lake Kumba, Nyumba ya Mungu and Pangani Falls. Error bars represent 95% confidence intervals.

The key results of this study are that *O. niloticus* demonstrates faster growth relative to the indigenous *O. jipe*, but also that extent of differences varies among locations. Such differences may have multiple explanations. Since *Oreochromis* can respond rapidly to selection on body size traits (Hulata et al. 1986), and recent work has identified significant genetic differences in neutral markers among the three sampled *O. niloticus* populations (Shechonge et al. 2019a), then genetic variation may underpin growth differences among the populations of both species. This may reflect historic selection from impoundments or aquaculture facilities prior to being introduced, or fisheries-induced evolution (Heino et al. 2015). Alternatively, the different sampled environments may differentially favour the species, with conditions within the Nyumba ya Mungu reservoir particularly well suited to growth of *O. niloticus* relative to *O. jipe*. It is unknown to what extent these species use different niches within each of the sampled environments. To fully understand the underlying reasons for the differences in growth rates between and within species would require more detailed study of growth rates in common-garden conditions, in addition to an improved understanding of the relative differences among populations in habitat, diet and levels of fisheries exploitation.

Although our analysis of scale increments is suggestive of more rapid growth of *O. niloticus* than *O. jipe* in some circumstances, other phenotypic characters relevant to fisheries need to be assessed, including maximum length, age of maturity and food conversion rate. Higher individual growth rate need not translate into greater rate of total fish biomass production which is likely to be more relevant for small-scale fishery yields. Whether the observed differences will have relevance for ecological interactions between the species is also unclear. It is possible that a faster growth rate for the non-native *O. niloticus* may be advantageous when competing with *O. jipe* for limited resources, including food, breeding space or shelter from predators. This is potentially of concern given the Critically Endangered IUCN red list status of the *O. jipe*, linked to its narrow geographic range and overall decreasing population trajectory (Bayona and Hanssens 2006). In Lake Kariba, *O. niloticus* has been shown to possess faster growth rate than indigenous *Oreochromis mortimeri* (Trevawas, 1966; Chifamba and Videler 2014). This, coupled with evidence of a rapid population expansion of *O. niloticus* matching a decline in *O. mortimeri* from the late 1990s onwards (Chifamba 2006), and evidence of overlapping resource use patterns (Mhlanga 2000), is suggestive of the potential for *O. niloticus* to outcompete indigenous species. Equivalent monitoring of the abundance changes, resource use patterns and detailed analyses of life history parameters of *O. niloticus* and sympatric indigenous tilapia would help to understand the full effects of introductions of *O. niloticus* across East Africa.

**Table 1.** Number and average standard length (± standard deviation) of *O. niloticus* and *O. jipe* sampled from the three study locations.

|  | Lake Kumba |  | Nyumba ya Mungu |  | Pangani Falls |  |
| --- | --- | --- | --- | --- | --- | --- |
| | n | SL ± SD | n | SL ± SD | n | SL ± SD |
| *O. niloticus* | 71 | 10.88 ± 2.59 | 15 | 12.97 ± 4.58 | 26 | 6.51 ± 0.96 |
| *O. jipe* | 13 | 9.41 ± 1.31 | 18 | 11.80 ± 3.33 | 6 | 5.66 ± 0.65 |

## Acknowledgements

The work was funded by Royal Society-Leverhulme Trust Africa Awards AA100023 and AA130107. We thank the Tanzania Commission for Science and Technology (COSTECH) for research permits, and staff of the Tanzania Fisheries Research Institute for contributions to fieldwork.

